# The molecular architecture of vimentin filaments

**DOI:** 10.1101/2021.07.15.452584

**Authors:** Matthias Eibauer, Miriam S. Weber, Yagmur Turgay, Suganya Sivagurunathan, Robert D. Goldman, Ohad Medalia

## Abstract

Intermediate filaments are integral components of the cytoskeleton in metazoan cells. Due to their specific viscoelastic properties they are principal contributors to flexibility and tear strength of cells and tissues. Vimentin, an intermediate filament protein expressed in fibroblasts and endothelial cells, assembles into ~11 nm thick biopolymers, that are involved in a wide variety of cellular functions in health and disease. Here, we reveal the structure of in-situ polymerized vimentin filaments to a subnanometer resolution by applying cryo-electron tomography to mouse embryonic fibroblasts grown on electron microscopy grids. We show that vimentin filaments are tube-like assemblies with a well-defined helical symmetry. Their structure is comprised of five octameric, spring-like protofibrils harboring 40 vimentin polypeptide chains in cross-section. The protofibrils are connected by the intrinsically disordered head and helix 1A domains of vimentin. Individual filaments display two polymerization states characterized by either the presence or absence of a luminal density along the helical axis. The structure of vimentin filaments unveils the generic building plan of the intermediate filament superfamily in molecular details.

## INTRODUCTION

Vimentin is an intermediate filament (IF) protein which assembles into extensive cytoskeletal networks mainly in cells of mesenchymal origin ^1,2^. Vimentin IFs (VIFs) participate in a broad range of cellular processes, such as the development of focal adhesions ^3^, the protrusion of lamellipodia ^4^, the assembly of stress-fibres ^5^, the elongation of invadopodia ^6^, the regulation of the dynamic properties of microtubules ^7,8^, and even virus infection ^9,10^. Vimentin expression and the assembly of VIFs are canonical markers and regulators of the epithelial-mesenchymal transition ^11^ and are involved in many aspects of cancer initiation and progression ^12,13^, and other pathophysiological conditions ^14–17^.

More specifically, VIFs are essential components of cellular architecture with respect to the establishment and maintenance of the shape and mechanical integrity of cells ^18–20^. Furthermore, VIFs form highly dynamic cytoplasmic meshworks that extend throughout the cell and which can rapidly respond to changes in the cellular environment. This fast and versatile remodelling of the VIFs comprising these meshworks depends both on their ability to exchange subunits along the filaments ^21,22^, and on posttranslational modifications ^23,24^. Oxidative and electrophilic modifications of vimentin induce dramatic alterations in VIF architecture and meshwork organization ^25,26^.

Similar to other types of IF proteins ^27^, vimentin contains an elongated, central rod domain composed of four coiled-coil α-helical segments, termed 1A, 1B, 2A, and 2B, respectively, that are connected by flexible linkers. In turn, the rod domain is flanked by intrinsically disordered N-terminal head (H) and C-terminal tail (T) domains (Fig. 1a). Vimentin monomers assemble into parallel homodimers ^28^, ~49 nm in length ^29^. Snapshots into vimentin assembly in-vitro suggested that two dimers interact to form an antiparallel tetramer ^30^, with a length of ~61 nm and a molecular mass of ~214 kDa ^27,29^. This tetramer is the basic building block for the subsequent assembly of VIFs ^22,31^.

**Figure 1.**
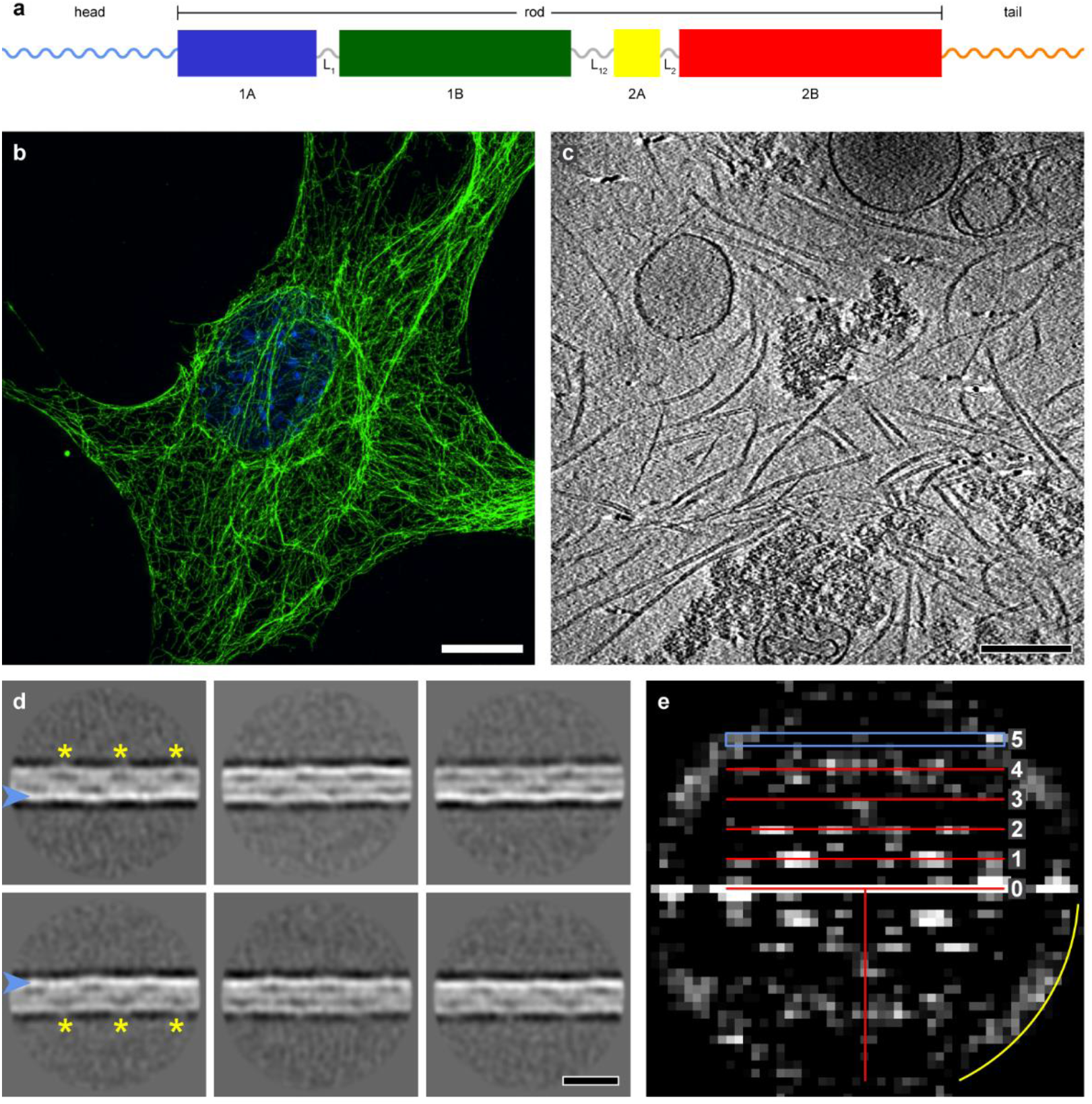
Cryo-ET and power spectrum analysis of VIFs. (**a**) Domain organization of a vimentin monomer. The central rod consists of predominantly alpha-helical domains (1A, 1B, 2A, 2B), connected by flexible linkers (L1, L12, L2). The rod is flanked by intrinsically disordered head and tail domains. (**b**) Slice of a 3D-SIM image of a MEF, fixed and stained with anti-vimentin (green) and the nucleus is stained with DAPI (blue). The VIF network extends over the whole cellular volume, with regions of lower and higher network density, and forms a cage around the nucleus. Scale bar is 10 µm. (**c**) Slice through a cryo-tomogram of a detergent treated MEF. The VIFs can be clearly detected. They consistently show no unravelling over the whole dataset. (**d**) The VIF state-1 class averages show a helical pattern with a repeat distance of ~180 Å (yellow asterisks). One side of the filament boundary appears pronounced in projection (blue arrowheads). Scale bar is 180 Å. (**e**) Combined power spectrum of the VIF state-1 class averages. The presence of layer lines confirms the helical architecture. The first layer line appears at ~1/186 Å and reflects the repeating pattern observed in the class averages. The layer line and peak distribution with a meridional reflection on the fifth layer line at ~1/37 Å is compatible with a helical assembly of five subunits per repeat. The yellow arc in the power spectrum indicates 1/26 Å.

The next steps in the hierarchy of vimentin polymerization involve the lateral and longitudinal concatenation of tetramers into higher order structures that contain multiples of 4 vimentin polypeptide chains. These higher order structures assemble into protofilaments (tetramers attach longitudinally, 4 monomers in cross-section) ^27^, protofibrils (8 monomers in cross-section) ^32^, and forming unit-length filaments (ULFs) ^33–36^, structures which are highly variable regarding their molecular mass (32, 40, or 48 monomers in cross-section). Ultimately, ULFs fuse longitudinally to form mature VIFs ^31,37,38^.

The extensive structural polymorphism of VIFs together with their elongated, flexible building blocks and intrinsically disordered domains, impose major challenges for the determination of their 3D structure. Nevertheless, a divide and conquer X-ray crystallographic analysis of overlapping vimentin fragments have provided an atomistic model of the nearly complete rod domain ^39^. Based on molecular dynamic simulations much of the structure of the vimentin tetramer has been determined ^29,40,41^. Despite this progress, the structure of VIFs in their fully assembled filamentous state within cells is required for a deeper understanding of the unique properties and many functions of VIFs.

Here, we have determined the structure of in-vivo polymerized VIFs employing cryo-electron tomography (cryo-ET) and helical reconstruction procedures. Through the classification of filament segments and the assembly of class averages into filaments of substantial length, we have been able to uncover the polymorphism along the VIFs. Disentangling individual filament states, a 3D reconstruction of VIFs has been achieved providing subnanometer resolution.

VIFs feature a helical assembly of 5 spring-like protofibrils built from the 1B, 2A, and 2B domains of vimentin and the protofibrils are linked together by the 1A and H domains. Our analysis provides detailed insights into the architecture of VIFs and reveals the role of individual vimentin domains within the filament structure.

## RESULTS

### Cryo-ET of VIFs

Mouse embryonic fibroblasts (MEFs) are widely used for investigating the cellular organization and function of VIFs. In this study we used MEFs that form a typical VIF network as revealed by 3D structured illumination microscopy (3D-SIM) imaging (Fig. 1b). The cells were cultured on electron microscopy grids and subjected to a permeabilization procedure that preserves the VIF network, while removing soluble components from the cytoplasm and the nucleus ^42^. The VIF networks in detergent treated cells closely resemble those of intact cells (Supplementary Fig. 1a).

Using these MEF preparations, we acquired 225 cryo-tomograms in areas with varying numbers of VIFs (Fig. 1c, Supplementary Fig. 1b, Supplementary Movie 1). The VIFs exhibit a tube-like appearance with no indications of partial unravelling, suggesting a reduced heterogeneity in comparison to the results obtained with bacterial expressed vimentin assembled into VIFs in-vitro ^43^. Using a convolutional neural network ^44^, we identified and extracted 390,297 segments of VIFs with a box size of 65 × 65 × 65 nm^3^. These segments were projected along the direction of the electron beam and subsequently subjected to a 2D classification procedure ^45,46^ (Supplementary Fig. 1c). Surprisingly, this classification revealed two distinct structural patterns in VIFs (Supplementary Fig. 2); one is helical-like and the other is rope-like in appearance. We term these VIF state-1 and VIF state-2, respectively.

### Repeat distance and power spectrum analysis

VIF state-1 class averages (Fig. 1d, Supplementary Fig. 3a) reveal two tracks of a repeating pattern of elongated, low-density regions, which are displaced relative to each other, run parallel to the outer boundaries of the filament, and are accompanied by local increases of curvature and outward protrusions (Fig. 1d, yellow asterisks). The repeat distance of the pattern is ~180 Å, as measured by autocorrelation analysis ^47^ (Supplementary Fig. 3b). Additionally, the class averages reveal that one filament wall appears more pronounced in projection than its counterpart (Fig. 1d, blue arrowheads), which suggests an odd number of subfilamentous entities building the filaments (Supplementary Fig. 3c).

Subsequently, we combined all state-1 class averages into a common power spectrum, which shows helical layer lines (Fig. 1e), thereby confirming the helical symmetry of state-1 VIFs. A clearly detectable layer line is positioned at ~1/186 Å, which corroborates the previous autocorrelation measurement. Moreover, the Bessel peak distribution along the layer line spectrum is compatible with a helical assembly that packs five building blocks in one helical pitch, which would correspond to the observed asymmetry in the class averages. Consequently, on the fifth layer line, which is positioned at ~1/37 Å (Fig. 1e, blue rectangle), a meridional peak can be detected.

### Determining the helical symmetry of VIFs

We confirmed the helical indexing scheme by an exhaustive helical parameter search on both filament states. Therefore, we merged all vimentin segments into one stack of particles (133,780 segments of size 65 × 65 nm^2^) and performed an initial helical 3D classification. In order to understand the basic diameter and low-resolution features of the six class averages obtained, all helical information was discarded in the first instance and the 3D class averages were fully symmetrized along their central axis.

Interestingly, the class averages differ in the presence of a luminal density along their central axis (Supplementary Figs. 4a&b). Determinations of their diameter suggest that the increases in the diameter of VIFs coincide with increases in luminal density (Supplementary Fig. 4c). The maximum diameter difference detected is ~2 nm. The outer tube diameter is 12.7 nm ± 0.7 nm and the centre-to-centre distance between the tube walls is 9.4 nm ± 0.8 nm with a tube wall thickness of 3.3 nm ± 0.4 nm. Based on these measurements, we define VIF state-1 as those polymers exhibiting a pronounced luminal density and VIF state-2 missing the distinct mass along their central axis.

In the following, we examined whether the two polymer states are similar with respect to their helical symmetry, so that their primary structural difference would be caused by the luminal density. Therefore, we used the particle sets of the six class averages from before as separate inputs for a series of unbiased, helical 3D refinements ^48,49^, assuming that the true helical symmetry generates optimal resolution. Thereby, we searched through a wide spectrum of helical rise values from 30 Å to 69 Å at a constant helical pitch of 186 Å. This brute-force search visited a wide range of helical geometries that contain between 2.7 to 6.2 asymmetric units in the helical pitch length. The results show that four classes reached optimal resolution values around a helical rise of 37 Å (Supplementary Fig. 4d), and the mean resolution over all classes is optimal at a helical rise of ~37 Å (Supplementary Fig. 4e). Subsequently, we applied the helical symmetry obtained for 3D classification of the vimentin segments and confirmed that the features of the reconstructed class averages in projection match the experimental 2D class averages (Supplementary Fig. 4f).

The results of our analyses reveal that the helical symmetries of VIF state-1 and VIF state-2 are similar to a first approximation, indicating therefore the luminal density does not significantly alter the helical symmetry of the surrounding filament tube. In order to assess the validity of the helical rise measurements, we assumed a closely-packed helical structure, so that the helical rise could be estimated based on the helical pitch and the filament diameter ^50^. Here, a helical pitch of 186 Å and a filament diameter of 12.7 nm results in a helical rise of 36.8 Å, and the maximum diameter difference of ~2 nm between the two polymer states is implying a variation in the helical rise between 36.1 Å and 37.5 Å. This interval defines the theoretical difference between VIF state-1 and VIF state-2 in terms of the helical rise, that is compatible with the measured diameter polymorphism of VIFs (Supplementary Fig. 4c). Three independent sets of measurements consistently reveal a helical pitch of ~186 Å and a helical rise of ~37 Å. These values are in agreement with cross-linking experiments suggesting a helical rise of 35.5 Å for VIFs (z_a_value in ^51^), as well as scanning transmission electron microscopy mass-per-length measurements. Assuming that the vimentin tetramer is the asymmetric unit of the helical assembly ^22,31^, a mass-per-length of 56 kDa/nm (third peak in Fig. 9a in ^33^) directly ^52^ yields a helical rise of 38.3 Å.

### Long-range tracing of VIFs

Resolving the structural fluctuations along the VIFs requires the reconstruction of extended polymer stretches ^45,53^. Therefore, we extracted smaller segments (38 × 38 nm^2^) and decreased the distance separating them, in order to increase the number of samples to more accurately track the natural bending of the VIFs. After several rounds of 2D classification, the initially selected ~1.1 million particles were concentrated to 615,106 segments and grouped into 100 class averages (Supplementary Fig. 5). These class averages were then transformed, so that their position and orientation matched the corresponding raw segments, and by this means the VIFs were assembled from the class averages. As a result, their signal-to-noise ratio is greatly improved compared to the raw images of VIFs. An example of an assembled VIF is shown in Fig. 2a. The change in structural patterns along a single VIF can be observed over a substantial length. To complete the long-range tracing of VIFs, we applied a straightening procedure to unbend the filaments ^1,54^.

**Figure 2.**
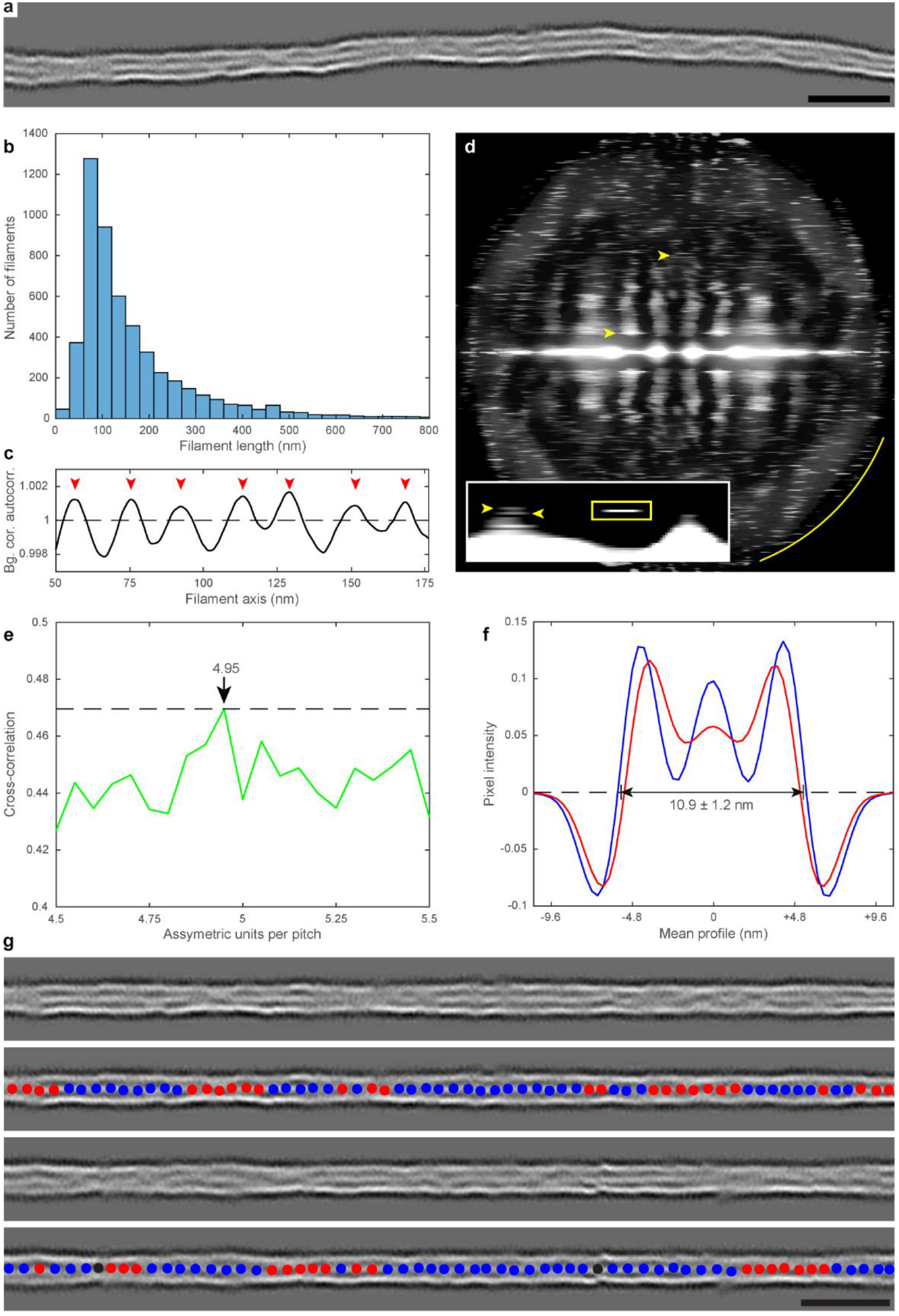
Long-range structural patterns of VIFs. (**a**) Filament assembly allows to follow the progression of the VIFs with improved signal-to-noise ratio over a substantial length. Scale bar is 35 nm. (**b**) Length distribution of the assembled filaments. (**c**) The long-range repeat distance of the VIFs was measured. The plot shows the autocorrelation signal after background correction. The mean distance between the red arrowheads is 186.5 Å ± 26.0 Å. (**d**) The assembled VIFs were combined into one power spectrum. A dense pattern of layer lines can be detected, reflecting the diffraction pattern of a discontinuous helix. The final structure of state-1 VIFs converged to a helical pitch of 185.8 Å and a helical rise of 37.1 Å. At both positions sharp layer lines can be detected in the power spectrum (yellow arrowheads). The yellow arc indicates 1/16 Å. In the inset at the putative pitch region a sharp layer line (yellow rectangle) can be detected at 185.6 Å ± 4.9 Å. It can be observed that the layer lines split into triplets (yellow arrowheads). (**e**) The splitting of the layer lines was modelled. At a packing of 4.95 ± 0.05 vimentin tetramers in the length of a helical pitch the similarity between the experimental power spectrum and the simulated power spectra is maximized, thereby confirming the helical indexing scheme based on long-range structural patterns of VIFs. (**f**) Two radial profiles along VIFs in state-1 and state-2 are plotted (blue and red curves, respectively). In state-1 the luminal density is almost as dense as the filament walls. In state-2 the luminal density appears suppressed and the filament walls move inwards. (**g**) Two assembled VIFs are shown. The blue and red dots along the filament axis indicate the position of state-1 and state-2 segments, respectively. Scale bar is 35 nm.

In total, we assembled 5205 VIFs of different lengths (Fig. 2b). We selected a subgroup of 389 of these VIFs with a length of ≥353 nm. These were boxed to an equal length (1024 pixel) and we measured the repeat distance along these VIFs employing autocorrelation (Fig. 2c). The result showed a long-range repeat distance of 186.5 Å ± 26.0 Å, supporting the previous measurements on single class averages.

Next, we calculated a combined power spectrum of these VIFs (Fig. 2d). Due to the increased resolution, a dense spectrum of layer lines is revealed and Bessel peaks confirming the previously measured helical pitch and rise values can be clearly detected (Fig. 2d, yellow arrowheads). The inset in Fig. 2d focuses on the most significant Bessel peaks above the background, in a region around the putative helical pitch. The layer line at 185.6 Å ± 4.9 Å (yellow rectangle) confirms the previous helical pitch measurements and a subtle splitting of the layer lines can be observed (yellow arrowheads).

These results support the notion that the helical pitch of VIFs is determined by a characteristic, constant length within the vimentin tetramer. Indeed, it was shown based on molecular dynamics simulation ^29^ and cross-linking experiments ^55^ that the lateral stagger of the tetramer in the A_11_ binding mode (characterized by largely overlapping 1B domains) is similar to the helical pitch length which we have determined here. However, if the lateral spacing between the subfilamentous entities within the filaments can be modulated, for example due to post-translational modifications ^23^, the number of tetramers that can be packed into the constant helical pitch length will vary ^56^. This would result in fluctuations of both helical rise and filament diameter (Supplementary Fig. 4c), and ultimately creates a high density of layer lines in the power spectrum.

In order to substantiate this view of a fine-tuned molecular packing of VIFs, the atomic model of the vimentin tetramer ^29^ was placed in one of the previously generated fully symmetric averages (Supplementary Fig. 4a), and subsequently helical symmetrisation was applied. In this way, VIF models of defined length and helical symmetry were produced, converted into electron density maps, and then simulated power spectra were computed. In turn, the similarity between the simulated power spectra and the experimentally obtained power spectrum of VIFs (Fig. 2d) was measured by cross-correlation.

Based on this procedure we simulated filament geometries with a packing of 4.5 to 5.5 vimentin tetramers into a constant helical pitch length of 185.6 Å. The resulting cross-correlation curve showed that the similarity between the power spectra simulations and the experimentally derived power spectrum is maximized for a packing of 4.95 ± 0.05 tetramers (Fig. 2e). Consequently, the helical symmetry of VIFs in-situ is compatible with a fine-tuned molecular packing that converges to five tetramers in one helical pitch.

We also sought to determine the structural features and distribution of the two polymerization states along the VIFs. To this end, we conducted a helical 3D classification of the vimentin segments in four classes (Supplementary Fig. 6a). We assigned two classes with luminal density as VIF state-1 (324,386 particles) and two classes without this density as VIF state-2 (290,720 particles). Subsequently, we identified the respective segment positions along the assembled filaments, added a binary label that encodes the luminal density filling state, and calculated a score for each filament, reflecting its fraction of state-1 segments.

Based on this score, we calculated the two filament profiles shown in Fig. 2f. Here, we averaged the radial profiles of two sets of 300 assembled VIFs (length ≥22 nm), either with maximal or minimal luminal density score (blue and red profiles, respectively). The mean outer diameter of the VIFs in projection is 10.9 nm ± 1.2 nm. The blue profile indicates a pronounced luminal density in state-1 VIFs, whereas the red profile shows clearly that the luminal density in state-2 VIFs is reduced and their filament boundaries are more oriented towards the less dense filament lumen. The different structural motifs of the two states along the VIFs are clearly detected and it appears that segments in the same state form clusters along the VIFs (Fig. 2g, blue and red dots, respectively).

### Reconstruction of VIFs at subnanometer resolution

In order to determine the 3D structure of VIFs, we selected a class of 58,952 vimentin state-1 segments (Supplementary Figs. 6a-c) for further refinement. The helical parameters of the selected segments were validated by an exhaustive search (Supplementary Figs. 6d&e) and used as starting values for a 3D refinement procedure employing a local helical symmetry search using the RELION helical toolbox ^48^. The refinement procedure converged to a helical rise of h_f_=37.1 Å and a helical twist of t_f_=71.9°, which translates into n=5.0 asymmetric units packed into a helical pitch of 185.8 Å. The corresponding layer lines are annotated in Fig. 2d (yellow arrowheads).

The 3D structure of state-1 VIFs was resolved to 9.6 Å (Supplementary Fig. 7). The reconstruction is shown as a whole in Fig. 3a. A cut open view is presented in Fig. 3b containing the luminal density as it proceeds along the central axis of the filament. In cross-section the luminal density appears elliptical, with one axis of 3.7 nm ± 0.3 nm and the second axis of 3.1 nm ± 0.3 nm. Furthermore, the luminal density is oriented towards identical positions along the filament tube in steps of the helical pitch (Fig. 3b, squares).

**Figure 3.**
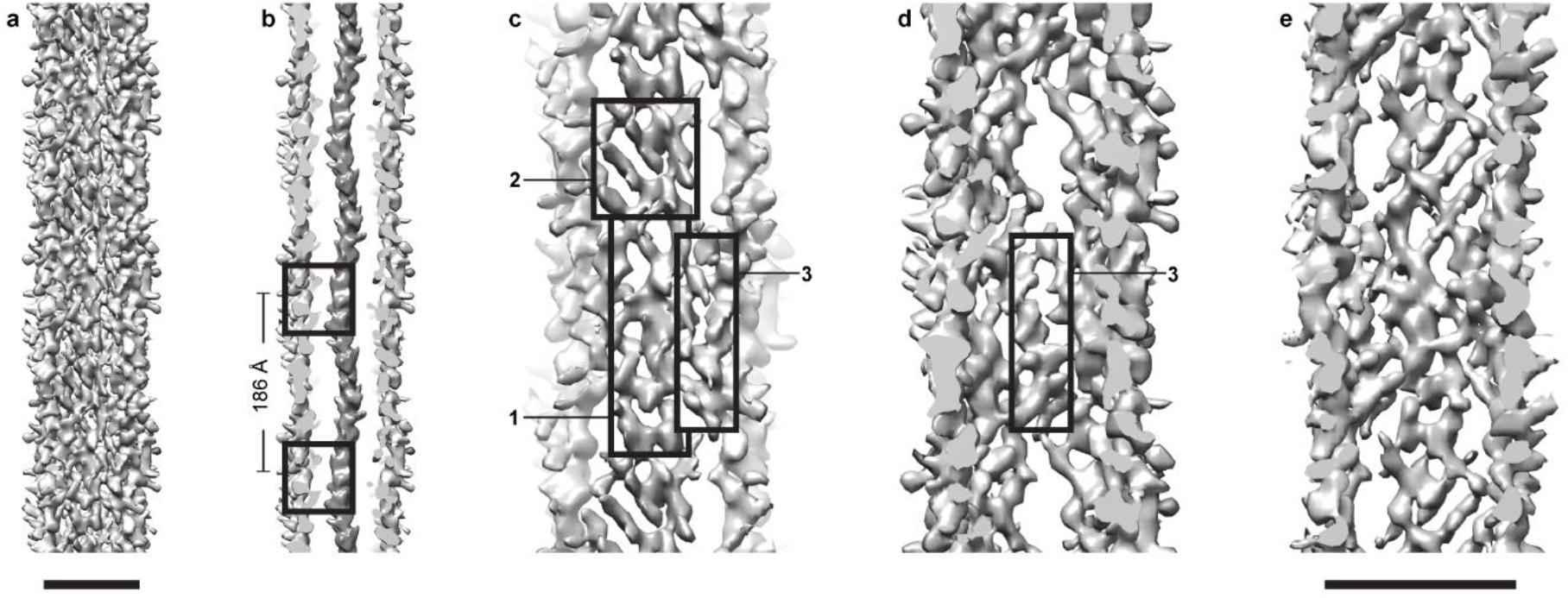
Helical 3D reconstruction of state-1 VIFs. (**a**) View of the complete structure of a vimentin state-1 polymer. Scale bar is 10 nm. (**b**) Cut open view of the VIF structure. The luminal density proceeds along the central axis and is oriented towards identical positions along the filament tube every ~186 Å, as indicated by the two squares. (**c**) Front view of one of the five protofibrils. The repeating unit can be subdivided into three characteristic regions. The first region (rectangle-1) exhibits a rod like structure, that runs parallel to the filament axis. The rod region is continuous with the triplet pattern of the hub region (rectangle-2). Densities that connect the protofibrils (the glue regions, rectangle-3) are staggered laterally relative to the rod and hub. (**d**) View from inside the VIF tube in one of the glue regions (rectangle-3). (**e**) View from inside the filament tube in the repeating unit of a protofibril. Scale bar is 10 nm.

Our analysis shows that in-situ polymerized VIFs are tube-like structures formed by 5 units, presumably protofibrils ^32^ composed of 2 tetramers in cross section (see below). In Fig. 3c we show three regions of a protofibril. The rod region (rectangle-1) is an elongated density along a protofibril that is delimited from the neighbouring protofibrils by openings in the structure. Along a protofibril, the rod region transitions into the hub region (rectangle-2), which can be identified by a characteristic triplet pattern. Interestingly, laterally staggered to the rod and hub regions, discrete densities that bridge between neighbouring protofibrils can be identified (rectangle-3). We propose that these regions serve as glue domains responsible for maintaining the structural integrity of VIFs by anchoring neighbouring protofibrils. Seen from inside of the VIF (Fig. 3d), the interactions of the protofibrils through the glue regions are apparent, and in Fig. 3e the complete repeating unit of a protofibril is shown.

In order to determine the molecular architecture of the VIFs, we constructed a model based on the measured helical symmetry and the atomic model of the vimentin tetramer ^29^. The consistency of the VIF model with the experimentally obtained structure was validated by docking, reaching a correlation coefficient of 0.81 (Supplementary Fig. 8, Supplementary Movie 2). The VIF model (Fig. 4, Supplementary Movie 3) reveals 8 vimentin polypeptide chains in cross-section of a protofibril and therefore 40 in cross-section of a filament ^36^. Another direct consequence of the helical symmetry which we have obtained is the mass-per-length of the VIF model (57.8 kDa/nm), which perfectly matches the data obtained by scanning transmission electron microscopy ^33,34^.

**Figure 4.**
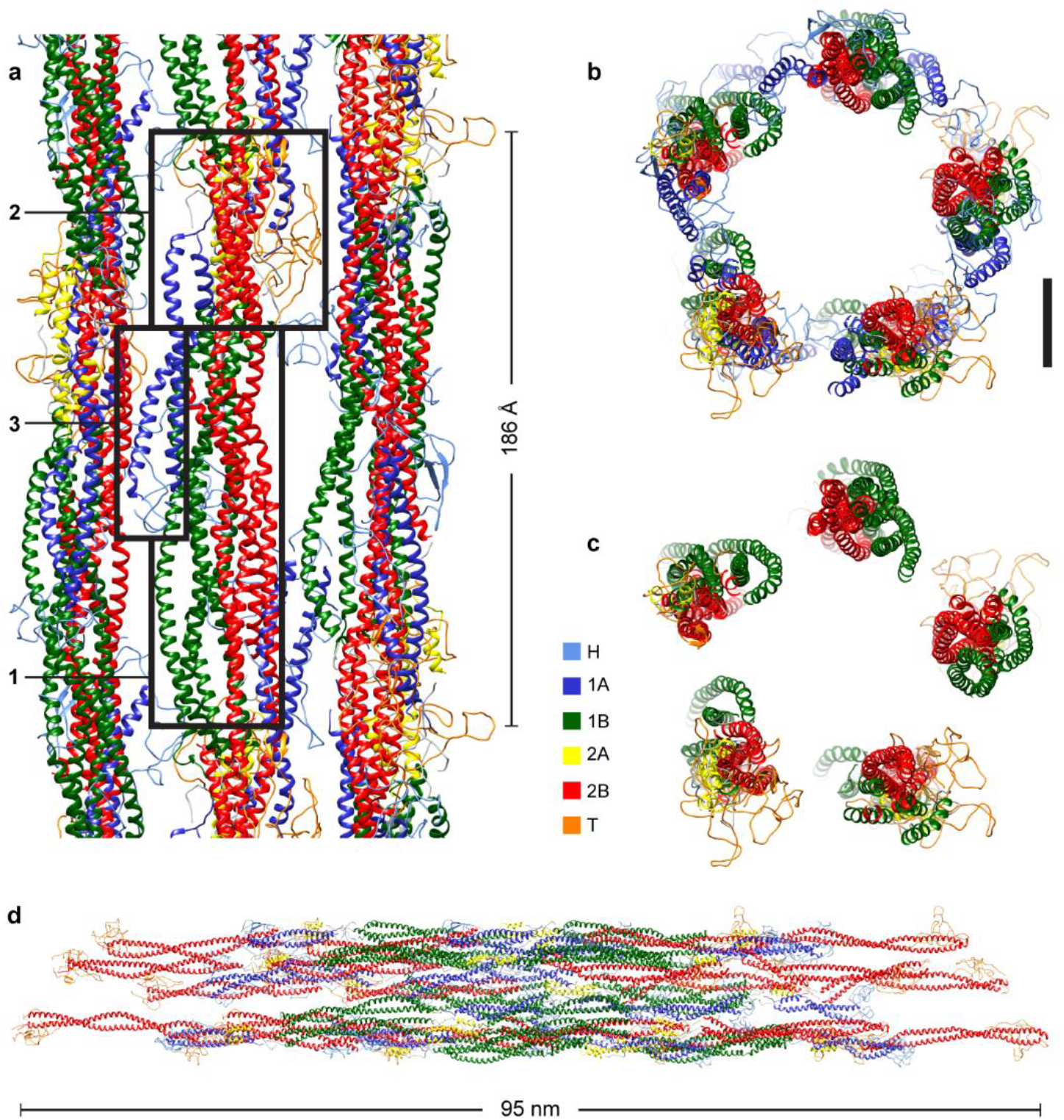
Molecular model of VIFs. (**a**) The VIF model indicates that the rod regions (rectangle-1) are assembled from parallel aligned 1B and 2B domains. In the hub regions (rectangle-2) the 2A domains are concentrated and the T domains are protruding away from the filament surface. Furthermore, from the hub regions the 1A domains extend into the space between the protofibrils where they form together with the H domains the glue regions (rectangle-3). The different protein domains are color-coded according to the color key. (**b**) Cross-section view of the VIF model. The five protofibrils are linked together by the 1A and H domains. Scale bar is 3 nm. (**c**) This becomes evident by omitting the 1A and H domains in the cross-section view. (**d**) Molecular model of an ULF. The 2B and T domains are accessible at both ends.

The model we have derived from our analyses shows that the rod region of a protofibril is assembled by interactions between the 1B and 2B domains of adjacent vimentin protein chains (Fig. 4a, rectangle-1). Further along a protofibril, the 2B domains enter the hub regions (Fig. 4a, rectangle-2), where the 2A domains are concentrated and T domains are protruding from the surface of the VIFs, explaining the protrusions previously detected in the class averages (Fig. 1d, yellow asterisks). Furthermore, the 1A domains extend from the hub regions into the space between the protofibrils. Together with the H domains the 1A domains form the glue regions (Fig. 4a, rectangle-3).

A cross section view of the VIF model is shown in Fig. 4b. This view highlights the essential role of the 1A and H domains for filament assembly, in agreement with the findings in previous studies ^18,28,33,57^. Clearly, the model shows that the 1A and H domains are the links between the spring-like protofibrils, as visualized by omitting the respective vimentin domains in the filament model (Fig. 4c).

A model of the vimentin ULF ^34^ is shown in Fig. 4d. Interestingly, both ends of the ULF are formed by the 2B and T domains of the vimentin protein, suggesting that these domains play a vital role in the elongation of VIFs. Indeed, a mimetic peptide that interferes with the 2B domain drastically alters the formation of ULFs and their longitudinal annealing, as demonstrated in a previous study ^28^. Assembly of a VIF with a cross section of 40 polypeptide chains requires at least two helix-turns, i.e. 10 tetramers. An alternative ULF model constructed from 5 tetramers and an elongated VIF model are presented in Supplementary Fig. 9.

## DISCUSSION

In this work we reveal the structure of VIFs. We have generated a molecular model of VIFs that maps the position and spatial relationships of the vimentin protein domains within the polymer and explains its general building principles (Supplementary Fig. 10). The precision of the VIF model largely depends on two factors, the accuracy of the measured helical symmetry and the fidelity of the vimentin tetramer model ^29,40^. Since the measured helical symmetry (h_f_=37.1 Å, t_f_=71.9°) provides a subnanometer resolved 3D reconstruction of the VIFs, and agrees with previous mass-per-length measurements ^33,34^ and results obtained from cross-linking experiments ^51,55^, our model permits a high precision position and spatial relationship determination of the vimentin protein domains building VIFs.

The observed molecular architecture of in-situ polymerized VIFs is in excellent agreement with previously published results. These include the established roles of the highly conserved 1A and 2B domains in VIF assembly and elongation ^18,28,33,57^, the contribution of the H domains to the lateral interactions between the protofibrils ^23^, and the number of polypeptide chains per cross-section ^36^. Since the VIF model predicts an alternating pattern of A_11_ and A_12_ tetramer binding interfaces along a protofibril, the VIF model explains the previously determined cross-links as well ^27,32,55^ (Supplementary Fig. 10b).

However, the subnanometer structural determination of the VIFs does not reveal their handedness. Since the sign of the helical twist angle is a priori undetermined and the map resolution is not sufficient to directly derive it, we set −t_f_, so that the protofibrils coil around the helical axis in a left-handed manner, in agreement with previous considerations based on structural hierarchy ^27^.

Interestingly, cellular VIFs form a homogenous assembly with respect to the number of polypeptide chains per cross-section (40 vimentin monomers), compared to in-vitro assembled VIF preparations, where the distribution is more heterogenous (32, 40, and 48 vimentin monomers) ^33,34,36^, suggesting that the cell selectively controls the polymerization of the VIFs. Furthermore, no regions containing unravelled VIFs were detected in the tomograms, as described for VIFs polymerized in-vitro from bacterial expressed vimentin ^43^.

Nevertheless, the VIFs polymerized within cells exhibit considerable polymorphism. Indeed, we observed that the VIFs exist in two major polymerization states, which are characterized by the presence of a luminal density in state-1 VIFs, and their absence or significant suppression in state-2 VIFs. Following the course of single VIFs demonstrate that transitions between these two states are common. As a consequence, the diameter of VIFs varies locally with a maximum difference of ~2 nm, which implies fluctuations in the helical rise between 36.1 Å and 37.5 Å, assuming a constant helical pitch. Although this variability cannot be neglected for 3D structural determination, it has only a minor influence on the general architecture of VIFs. Therefore, we suggest that the tube-like structures of state-1 and state-2 are similar, but the main difference is the luminal density. The molecular identity of the luminal density is still not clear, although, it is likely composed of vimentin protofilaments.

The VIF structures described in this analysis are likely to retain many, if not all of their in-vivo regulated post-translational modifications and therefore are in a near native state reflecting their in-vivo structural parameters. This ensures that we are studying the assembly and structure of physiologically relevant VIFs, without the requirement for the denaturing and renaturing procedures typically used in studies of VIF structure, and without the prerequisite to define the conditions for the polymerization in-vitro.

In the future, the strategy developed for this study can be applied to reveal the structure of VIFs in a variety of cellular states, and can be utilized as a starting point for the structural characterization of other members of the IF superfamily of cytoskeletal proteins. Larger datasets in conjunction with future developments in image processing will make it possible to resolve complete IF structures at atomic resolution.

## Acknowledgements

This work was funded by grants from the Swiss National Science Foundation (SNSF 31003A_179418) and the Mäxi Foundation to O.M. We thank the Center for Microscopy and Image Analysis at the University of Zurich.

## Author contributions

M. E. analyzed cryo-electron microscopy data and developed the methods. M. S. W. and Y. T. prepared samples and recorded cryo-electron microscopy data. S. S. prepared samples and recorded light microscopy data. M. E. together with R. D. G. and O. M. conceived the research and wrote the manuscript.

## METHODS

### Cell lines and cell culture

The MEFs were cultured in DMEM (Sigma-Aldrich, D5671) supplemented with 10% FCS (Sigma-Aldrich, F7524), 2 mM L-Glutamine (Sigma-Aldrich, G7513) and 100 μg/ml penicillin/streptomycin (Sigma-Aldrich, P0781), at 37°C and 5% CO_2_ in a humidified incubator.

### Light microscopy

Sub-confluent cultures of MEFs growing on #1.5 glass coverslips were fixed with 4% paraformaldehyde for 10 min at RT. The fixed cells were permeabilized with 0.1% Triton-X 100 for 10 min at RT and then incubated with chicken anti-vimentin (1:200, 919101, Biolegend, CA, USA) for 30 min in phosphate buffered saline (PBS) containing 5% normal goat serum. This was followed by staining with goat anti-chicken Alexa Fluor 488 (1:400, A-11039, Invitrogen, CA) and DAPI in phosphate buffered saline for 30 min. The stained cells were mounted with ProLong Glass Antifade Mountant (Life technologies, Carlsbad, CA, USA).

MEFs were seeded on #1.5 glass coverslips and the next day they were washed with PBS/2 mM MgCl_2_ for 5 s followed by incubation with PBS containing 0.1% Triton X-100, 10 mM MgCl_2_, 0.6 M KCl and protease inhibitors for 25 s at RT. The cells were rinsed with PBS/2 mM MgCl_2_ for 10 s and subsequently incubated with 2.5 units/µl Benzonase for 30 min at RT. After rinsing with PBS/2 mM MgCl_2_ the cells were fixed with 4% paraformaldehyde for 5 min at RT. The fixed cells were then stained with chicken anti-vimentin (1:200, 919101, Biolegend, USA) and rabbit anti-lamin A (to determine the location of the nucleus) in PBS containing 5% normal goat serum for 30 min at RT followed by incubation with goat anti-chicken and anti-rabbit secondary antibodies (1:400, A-11039, A-11011, Invitrogen, USA) for 30 min at RT. Following washing in PBS, the stained cells were mounted with Prolong Glass Antifade Mountant (Life technologies, Carlsbad, CA, USA).

3D-SIM imaging was carried out with a N-SIM Structured Illumination Super-resolution microscope system (Nikon, Tokyo, Japan) using an oil immersion objective lens (SR Apo TIRF100X, 1.49 NA, Nikon). For 3D SIM, 26 optical sections were imaged at 100 nm interval. The raw SIM images were reconstructed with the N-SIM module of Nikon Elements Advanced Research with the following parameters: illumination contrast, 1.00, high-resolution noise suppression, 0.75, and out-of-focus blur suppression, 0.25. Brightness and contrast were adjusted for image presentation.

### Cryo-ET sample preparation

MEFs were grown to ~80% confluency on glow-discharged holey carbon EM grids (R2/1, Au 200 mesh; Quantifoil, Jena, Germany) prior to preparation for cryo-ET analysis. Grids that showed a relatively homogenous distribution of cells were selected using fine tweezers, and then washed in PBS/2 mM MgCl_2_ for 5 s. The grids were treated for 20-40 s in pre-permeabilization buffer (PBS containing 0.1% Triton X-100, 10 mM MgCl_2_, 600 mM KCl and protease inhibitors) and then rinsed in PBS/2 mM MgCl_2_ for 10 s. In the next step, the grids were treated with Benzonase (2.5 units/µl in PBS/2 mM MgCl_2_; Millipore, Benzonase Nuclease HC, Purity >99%) for 30 min at RT. After washing the grids with PBS/2 mM MgCl_2_, a 3 µl drop of 10 nm fiducial gold markers (Aurion) was applied to the grids. For vitrification the grids were manually blotted for ~3 s from the reverse side and plunge frozen in liquid ethane.

### Cryo-ET

Tilt series acquisition was conducted using a Titan Krios transmission electron microscope (Thermo Fisher Scientific, Waltham, USA) equipped with a quantum energy filter and a K2 Summit direct electron detector (Gatan, Pleasanton, USA). The microscope was operated at 300 keV. Tilt series were collected at a nominal magnification of 42,000x and the slit width of the energy filter was set to 20 eV. Super-resolution movies were recorded within a tilt range from −60° to +60° with 2° increments using SerialEM ^58^. The image stacks were acquired at a frame rate of 5 fps with an electron flux of ~2.5 e^-^/pixel/s. The tilt series were recorded with a total dosage of ~125 e^-^/Å^2^ and within a nominal defocus range between −2 μm to −6 μm. The super-resolution image stacks were drift-corrected and 2x binned using MotionCorr ^59^, resulting in a pixel size of the tilt series of 3.44 Å. For each tilt series image the defocus was measured and the contrast transfer function (CTF) was corrected by phase-flipping. Then, from each tilt series a 4x binned overview tomogram was reconstructed. The CTF correction and overview tomogram reconstruction was performed using MATLAB scripts (MathWorks, Natick, USA) derived from the TOM toolbox ^60,61^.

### Image processing

Algorithms from EMAN2 ^44^ were employed for training a convolutional neural network capable of segmenting VIFs in the overview tomograms. The segmentations were manually checked and cleaned from obvious false positive VIF detections in Chimera ^62^. Based on scripts derived from the ActinPolarityToolbox ^45^ (APT), two sets of segment coordinates were extracted from the segmentations. In the first set (second set) the picking distance along the VIFs was set to 165 Å (55 Å), resulting in 390,297 (1,148,072) segment coordinates. Next, based on the segment coordinates two stacks of subtomograms were reconstructed from the CTF corrected tilt series with the TOM toolbox. The dimensions of the subtomograms were 65 x65 × 65 nm^2^ and 38 x38 × 38 nm^2^, respectively, and the pixel size was 3.44 Å. Subsequently, APT scripts were applied to project the subtomograms, using a projection thickness of 331 Å for the first set and 220 Å for the second set. The size of the subtomogram projections derived from the first and second coordinate sets were 65 × 65 nm^2^ and 38 × 38 nm^2^, respectively.

In the following, VIF segments were subjected to extensive, unsupervised 2D classifications in RELION ^46,63^ (Supplementary Fig. 1c, Supplementary Fig. 2a, and Supplementary Fig. 5) in order to sort out false positive and low signal-to-noise ratio segments. As a consequence, the first particle set (second particle set) was concentrated to 133,780 (615,106) VIF segments. The Markov chain analysis (Supplementary Fig. 2d) was based on 2D classification of the first particle set into 50 classes (Supplementary Fig. 2a). Employing APT scripts, the segments were connected to filaments and the sequences of the class averages along the filaments were analysed. For this purpose, a 50×50 transition matrix (T)_ij_ was calculated, with each matrix element ij indicating the number of class average transitions from class average i to class average j along the filaments. Next, (T)_ij_ was analysed with the MATLAB function *dtmc*.

The assembly of VIFs (Fig. 2a and Fig. 2g) was based on the 2D classification of the second particle set into 100 classes (Supplementary Fig. 5). For this purpose, the transformations calculated for each segment (namely their in-plane rotation angle psi and their xy-translation) were inverted and applied to the respective class averages, so that the inversely transformed class averages match position and orientation of the initial segments in the image frame of the tomograms ^53^. As a result of this operation the VIFs are represented by the class averages, which drastically improves their signal-to-noise ratio compared to the raw filaments. Subsequently, the assembled VIFs were unbent based on a MATLAB algorithm derived from the ImageJ ^64^ straighten function ^1,54^.

The autocorrelation profiles (Supplementary Fig. 3b, Fig. 2c) were calculated with the TOM toolbox function *tom_corr*, either calculating autocorrelation functions of class averages or assembled VIFs. The respective autocorrelation functions were averaged and displayed as profile plots. The power spectra (Fig. 1e, Fig. 2d) were calculated with the TOM toolbox function *tom_ps*, either calculating an averaged power spectrum from a series of class averages or a series of assembled VIFs. In the second case the assembled filaments were boxed to an equal length of 1024 pixel before calculating the averaged power spectrum.

The helical parameter searches (Supplementary Fig. 4d, Supplementary Figs. 6d&e) utilized a MATLAB script that created and executed RELION commands, and after completion analysed the results. Here, the RELION function *relion_refine* was used extensively in combination with the options –*auto_refine* (to ensure gold-standard resolution measurements ^49^) and –*helix* (to perform helical reconstruction ^48^). In order to scan a certain helical parameter interval, the options –*helical_rise_initial* and –*helical_twist_initial* were varied accordingly between the Refine3D jobs. Subsequently, the gold-standard resolution values were plotted and the helical parameter combination with the optimal resolution value was identified.

In order to create modelled power spectra of VIFs, the vimentin tetramer model, as obtained by molecular dynamics simulation ^29,40^, was fitted in the tube wall of one of the rotationally symmetrized class averages in Chimera (Supplementary Fig. 4a). Subsequently, defined helical parameter combinations were applied with the Chimera *sym* command to the fitted tetramer, thereby creating symmetry related tetramer copies. Next, the tetramers were converted into a 10 Å electron density map with the Chimera *molmap* command and the modelled electron density maps were elongated to 1024 pixel long filament models by using the RELION command *relion_helix_toolbox*. These structures were projected and power spectra were calculated. Based on the cross-correlation coefficient (TOM toolbox function *tom_ccc*), these modelled power spectra were compared with the experimentally measured power spectrum of VIFs (Fig. 2d). Subsequently, the cross-correlation values were plotted and the helical parameter combination with maximized similarity was identified (Fig. 2e).

Similarly, the VIF models (Fig. 4, Supplementary Fig. 9) were generated in Chimera. For this purpose, the vimentin tetramer model was fitted in the tube wall of the VIF structure (Fig. 3) and the measured helical symmetry (h_f_=37.1 Å, t_f_=71.9°) was applied with the Chimera *sym* command to the fitted tetramer, thereby creating a defined number of symmetry related tetramer copies (10 copies in Fig. 4d and 5, 10, and 30 copies in Supplementary Fig. 9). Subsequently, the tetramers were converted into an 8 Å electron density map with the Chimera *molmap* command and the modelled electron density map was docked into the VIF structure with the Chimera *fitmap* command (Supplementary Fig. 8).

The VIF helical lattice visualizations (Supplementary Figs. 10a&c) were created by transforming the modelled electron density map from cartesian to cylindrical coordinates with the TOM toolbox function *tom_cart2cyl*. This transformation was conducted for the densities of each of the vimentin protein domains individually. The transformed densities were projected and the resulting images were coloured and combined in one image.

### Visualization

All isosurface representations and the visualization of the VIF model were rendered with Chimera.

## Data availability

The algorithms developed in this work are available from the corresponding author upon request. The VIF structure is deposited in the Electron Microscopy Data Bank under the accession code EMD-13084.

## SUPPLEMENTARY MOVIES

**Supplementary Movie 1. Cryo-tomogram of a detergent treated MEF**. The movie shows successive slices of a representative cryo-tomogram of detergent treated MEFs. The field of view of the tomogram has a size of 1.41 × 1.41 µm^2^.

**Supplementary Movie 2. VIF model docked into VIF structure**. The movie shows the docking of the VIF model in the VIF structure (grey transparent density). The vimentin domains are color-coded according to the color key. The length of the filament shown is 60 nm.

**Supplementary Movie 3. VIF model rotating around the filament axis**. The vimentin domains of the rotating VIF model are color-coded according to the color key. The length of the modelled filament shown is 60 nm.

## SUPPLEMENTARY FIGURES

**Supplementary Figure 1.**
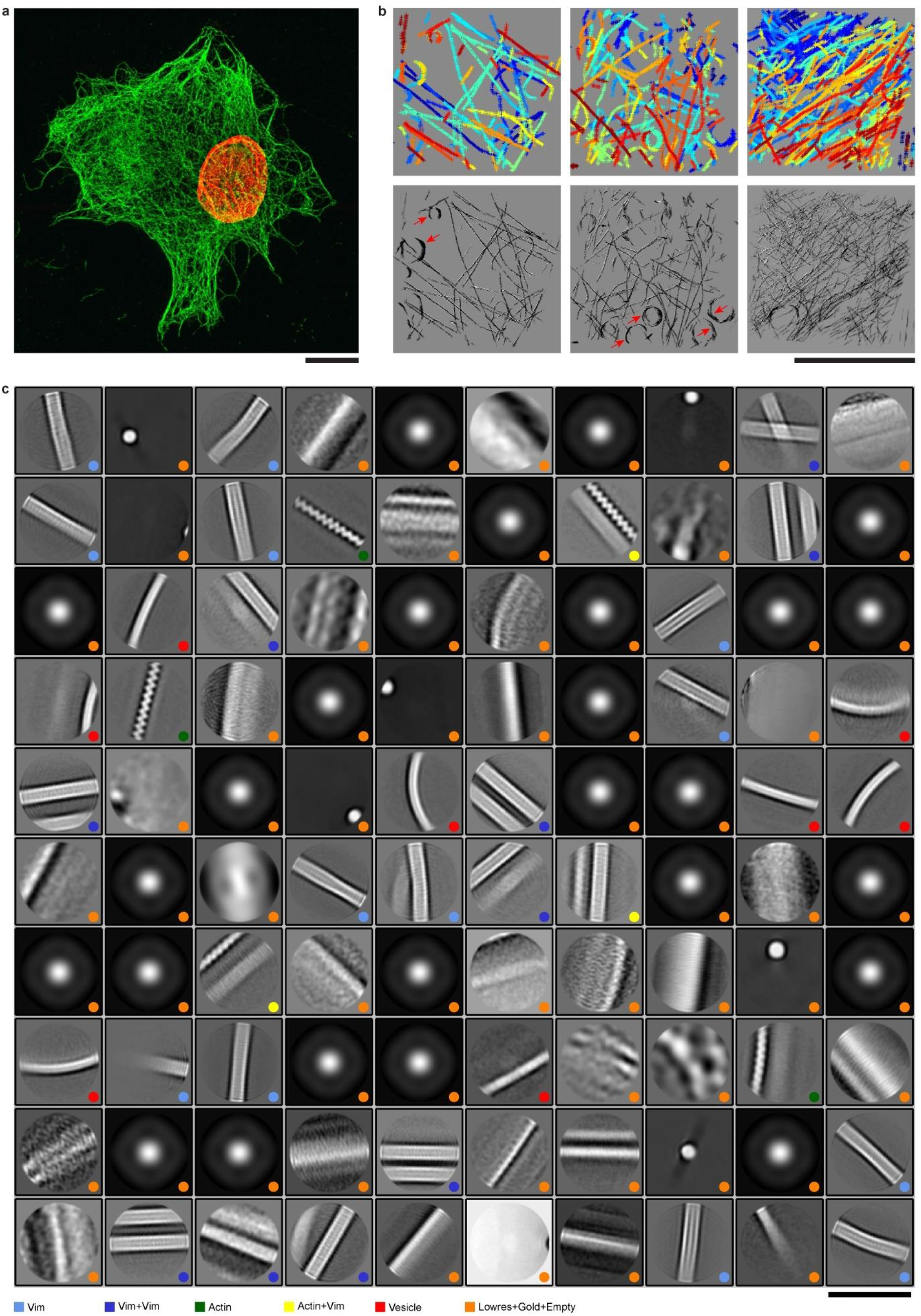
Deep classification of VIFs. (**a**) Maximum projection of a 3D-SIM image of a detergent treated MEF. The VIF network remains intact following the permeabilization procedure, fixation and staining with vimentin antibody (green). The cell nucleus is stained in red using lamin antibody. Scale bar 10 µm. (**b**) Three examples for picking VIFs with a convolutional neural network (top row). Because the filament detection is performed in 3D cryo-tomograms, VIFs from densely packed regions can be analyzed as well. Detected VIFs are shown in the lower panel. The convolutional neural network was over-picking membrane structures as VIFs (red arrows). Scale bar 1 µm. (**c**) Initial 2D classification of 390,297 projected vimentin segments (65 × 65 nm^2^, picking distance 165 Å, projection thickness 331 Å). The following patterns occur frequently in the class averages: Single VIFs (classes marked with light blue dots), two VIFs running parallel or crossing on top of each other (blue dots), single actin filaments (green dots), one VIF and one actin filament running parallel (yellow dots), and vesicle membranes (red dots). Other categories (very low-resolution class averages, gold markers or empty classes) are labelled with orange dots. Only particles from single VIF classes (light blue dots) were further analyzed. Scale bar 65 nm.

**Supplementary Figure 2.**
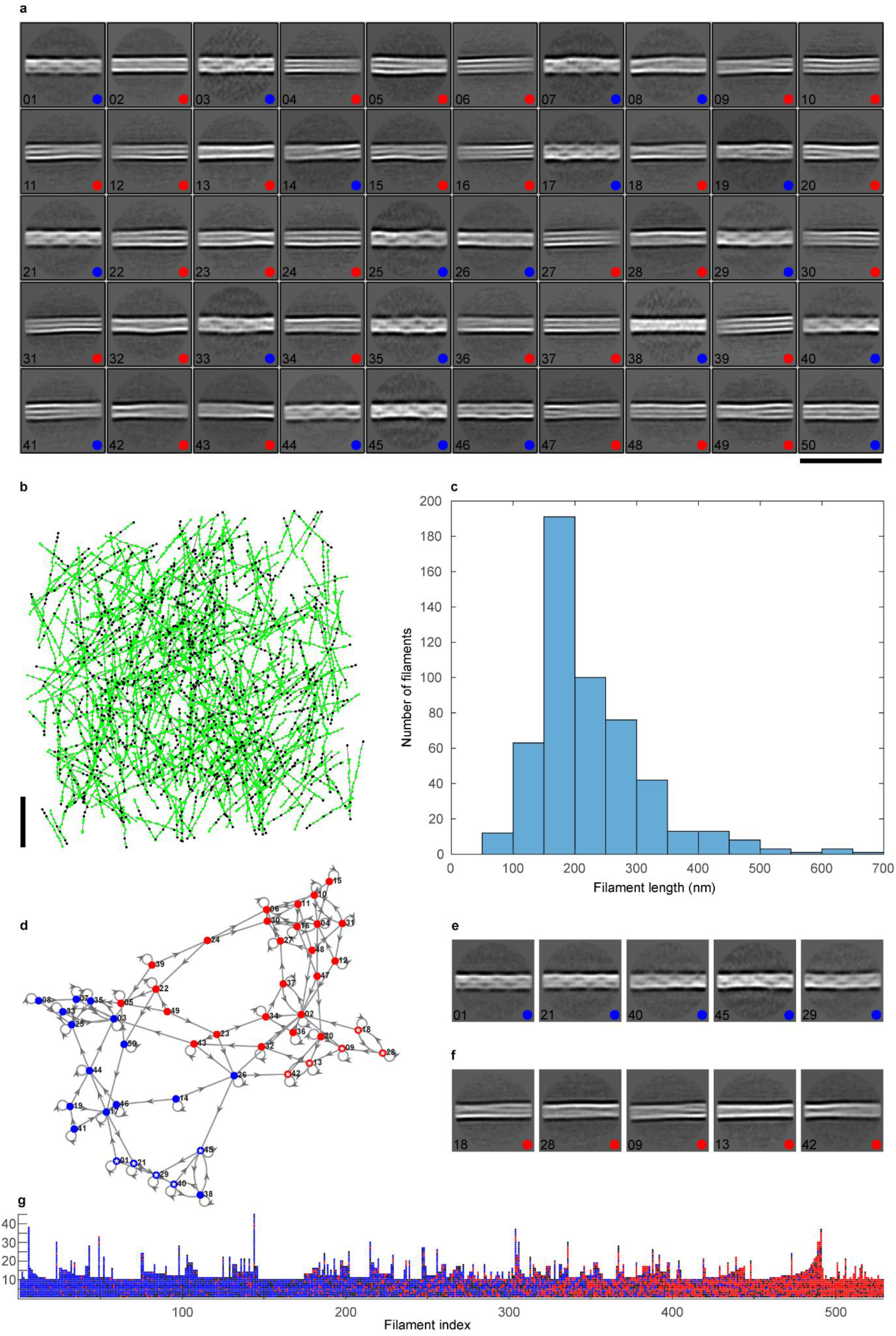
Extraction and pattern analysis of VIFs. (**a**) Class averages of VIFs split into two categories: The first group (VIF state-1) is characterized by similar helical patterns (class averages marked with blue dots). The second group (VIF state-2) is characterized by similar rope like patterns (class averages marked with red dots). The 2D classification is based on 133,780 segments. Scale bar 65 nm. (**b**) VIFs (in total 529 with a minimum length of 10 segments) were extracted based on the 2D classification shown in (**a**). Green dots represent the 3D coordinates of VIF segments, and segments plotted as black dots were sorted out during 2D classifications. Only filaments with a minimum proportion of 0.7 between processed segments (green dots) and total number of segments per filament were accepted. The orientations and positions of the filaments in the cryo-tomograms were randomly distributed. The z-coordinates of the segments were located within ±90 nm around the central xy-plane of the cryo-tomograms. (**c**) Length distribution of the extracted filaments. (**d**) The sequences of class averages were analyzed along the VIFs with a Markov chain analysis. State-1 class averages are closely connected (blue dots), as are state-2 class averages (red dots) as well. A sequence of state-1 averages (blue dots marked with white asterisks) is shown in (**e**), and a sequence of state-2 averages (red dots marked with white asterisks) is shown in (**f**). In (**g**) the extracted VIFs are plotted as columns, each segment is represented by a dot, and their colors indicate the structural pattern of the segments. Blue and red dots represent state-1 and state-2 structural patterns, respectively, and black dots are segments sorted out during 2D classifications. The VIFs are sorted from left to right based on their fraction of state-1 segments. VIFs which predominantly display state-1 or state-2 patterns can be observed, as well as filaments which are transitioning between these structural states.

**Supplementary Figure 3.**
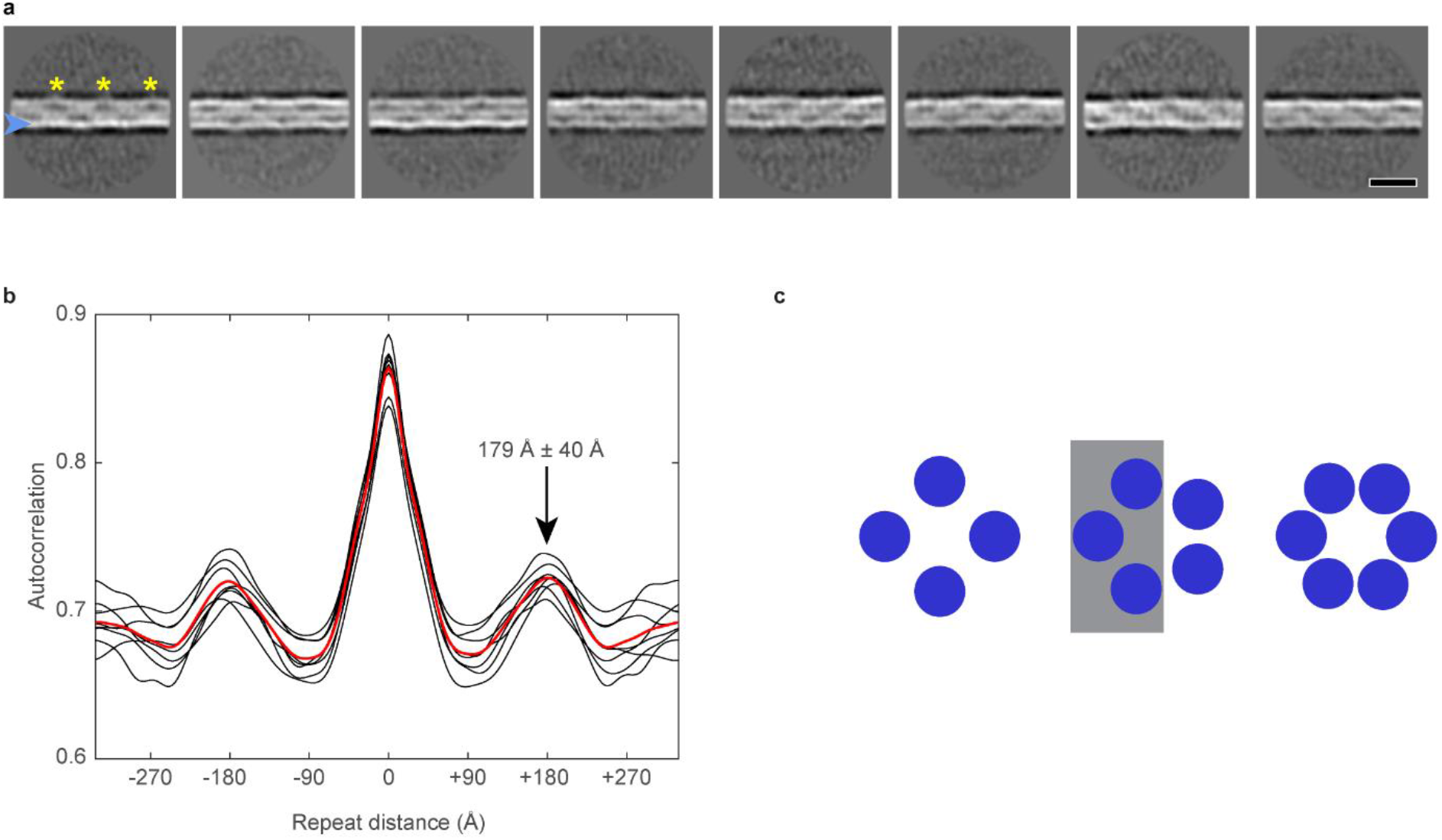
Repeat distance of VIF state-1 class averages. (**a**) Class averages showing wall asymmetry in projection (blue arrowhead) and repeating features like elongated, parallel low-density regions together with a local increase of boundary curvature and outward protrusions (yellow asterisks). Scale bar 180 Å. (**b**) The correlation of each class average with itself was calculated and the corresponding autocorrelation profiles along the x-axis were plotted (black lines). The red line is the averaged autocorrelation profile over all class averages. The pattern in the averages repeats at a distance of 179 Å ± 40 Å. (**c**) Model to explain the boundary asymmetry in the class averages (blue arrowhead in (**a**)). If the VIFs are assembled from five sub-filaments, one side of the VIF would appear brighter in projection (grey rectangle). However, if they are assembled from four or six sub-filaments the VIF boundaries would have similar densities in projection.

**Supplementary Figure 4.**
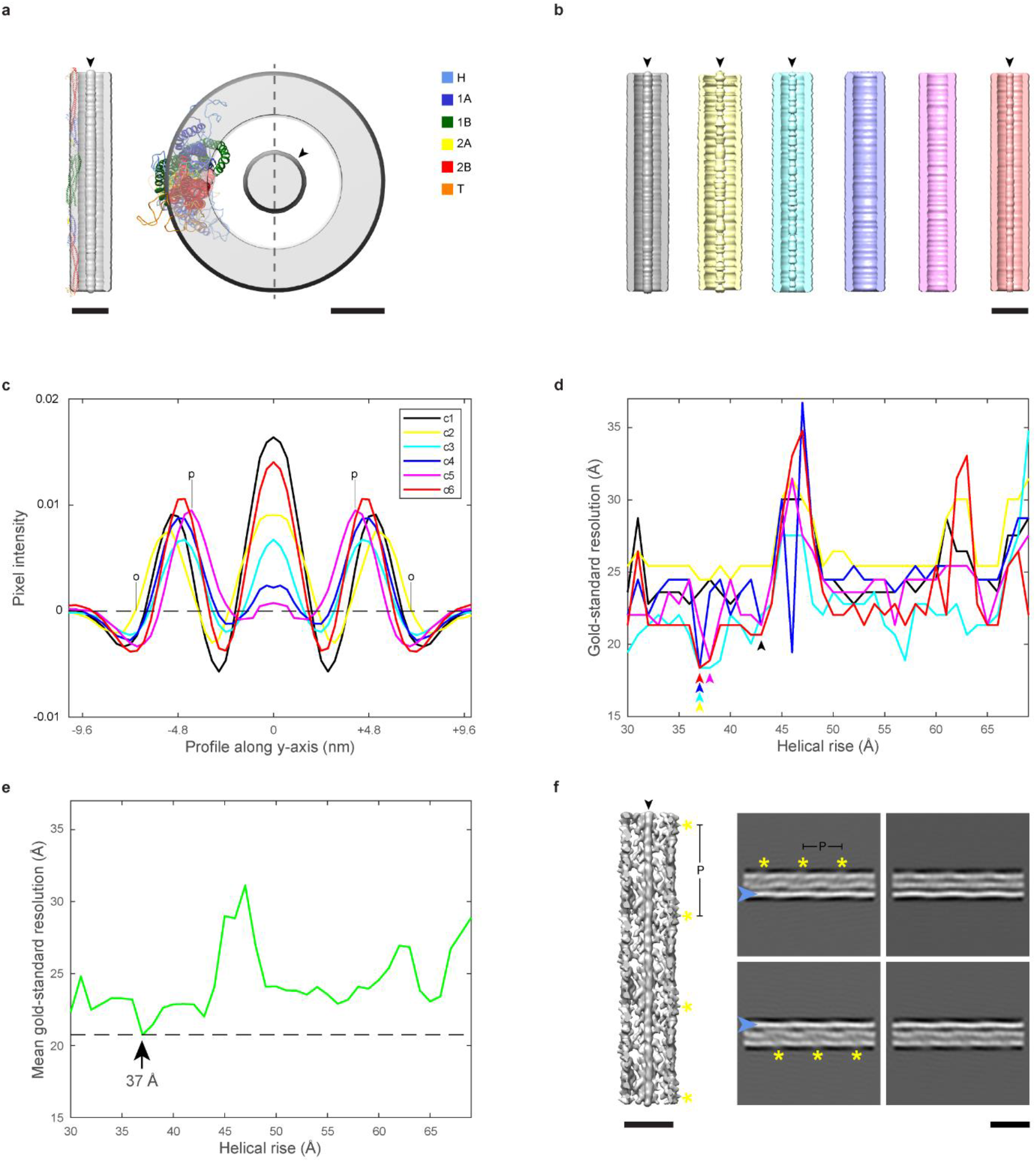
Helical parameter search. (**a**) VIF segments (133,780 particles of size 65 × 65 nm^2^) were classified in six classes and the resulting 3D averages were rotationally symmetrized along their central axis. One of the tubes is shown in a cut-open view on the left and shown from the top on the right. The black arrowheads indicate the luminal density. The vimentin tetramer model, as obtained by molecular dynamics simulation ^29^, has a length of ~61.3 nm and an average diameter in cross-section of ~3.6 nm. The tetramer model was fitted into the tube wall of the class average. Its tube wall thickness and curvature reflect the diameter and flexion of the tetramer model. The different domains of the tetramer are colored according to the color key. The vertical dashed line (y-axis) in the top view marks the path along which the profiles plotted in (**c**) were extracted. Left scale bar 10 nm and right scale bar 3 nm. (**b**) The symmetrized filament tubes are shown as cut-open views. The classification splits the particle set according to filament diameter and luminal density. Prominent luminal densities are marked with black arrowheads. Scale bar 10 nm. (**c**) Profile plots along the y-axis of the symmetrized class averages, referred to as c1-c6 in the legend. The average distance between the tube wall centers (p-p-distance, as exemplified for c5) is 9.4 nm ± 0.8 nm and the mean outer tube diameter (o-o-distance, as exemplified for c2) is 12.7 nm ± 0.7 nm. The average tube wall thickness is 3.3 nm ± 0.4 nm. (**d**) For each of the particle subsets, which are corresponding to the six symmetrized 3D class averages, 40 unbiased, gold-standard 3D refinements in RELION ^63^ were calculated with constant pitch of 186 Å, but varying the helical rise between 30 Å to 69 Å in 1 Å increments. The gold-standard resolution of the 3D refinements is plotted as a function of the helical rise. The global resolution optimum for the different particle subsets is indicated with arrowheads. (**e**) The green curve shows the average of the six resolution curves displayed in (**d**). The global resolution optimum is located at a helical rise of 37 Å. (**f**) The VIF structure shown here is the result of a helical 3D classification assuming a helical pitch of 186 Å and a helical rise of 37 Å. This class harbors a luminal density, which is marked with a black arrowhead. The filament wall exhibits outward protrusions, indicated with yellow asterisks, and their distance is equal to the helical pitch. Scale bar 10 nm. In addition, four reprojections of the filament structure are shown. The reprojections reproduce the features observed in the experimental class averages, that are the asymmetry of the VIF boundaries in projection (blue arrowheads) and parallel, low-density regions together with a local increase of boundary curvature and outward protrusions (yellow asterisks). Scale bar 180 Å.

**Supplementary Figure 5.**
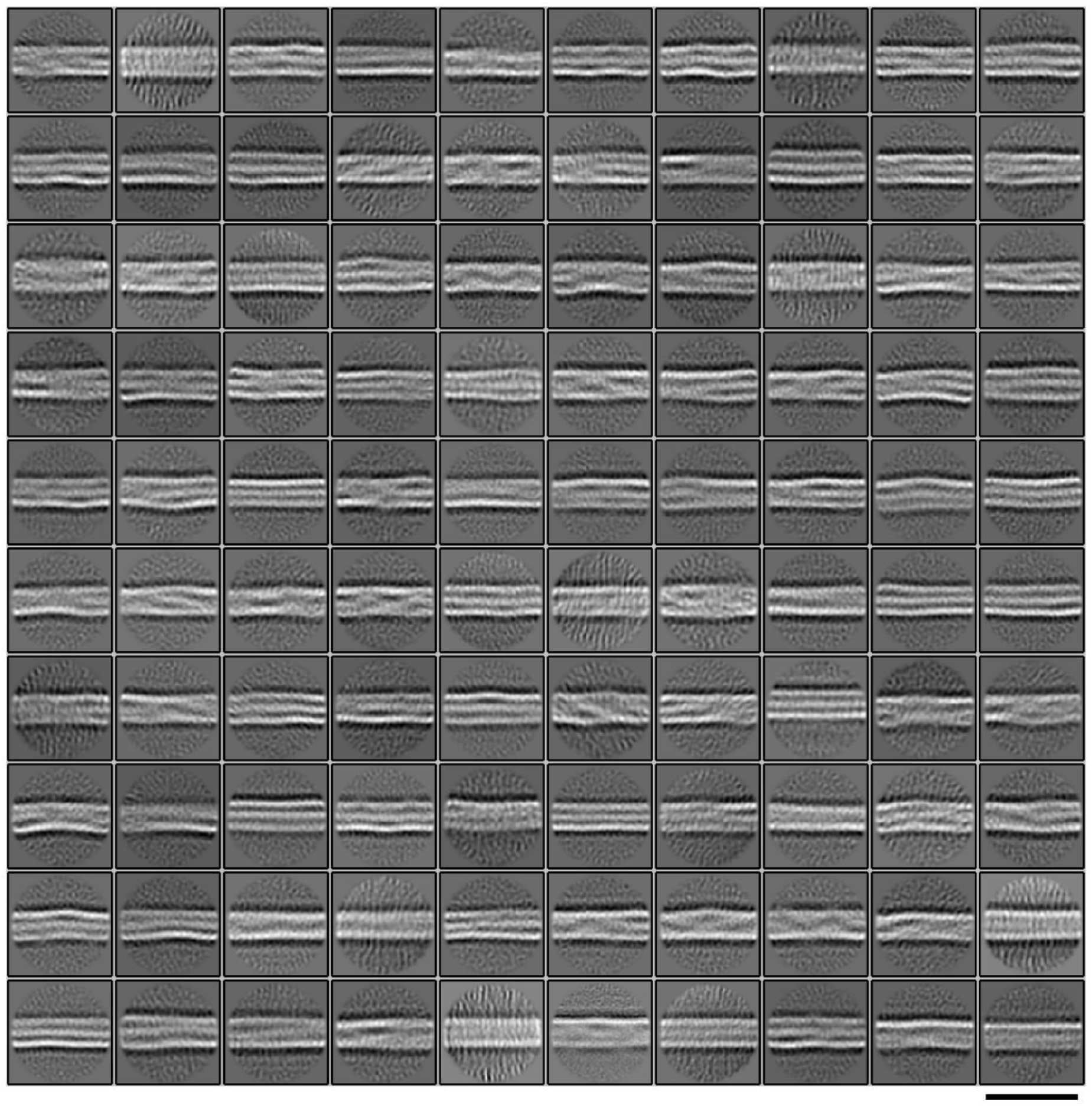
Extended 2D classification of vimentin segments. Vimentin segments (615,106 particles with a size of 38 × 38 nm^2^) were split by 2D classification in 100 classes. The picking distance between the segments was set to 55 Å and the projection thickness to 220 Å. The displayed class averages were used for subsequent filament assembly. Scale bar 35 nm.

**Supplementary Figure 6.**
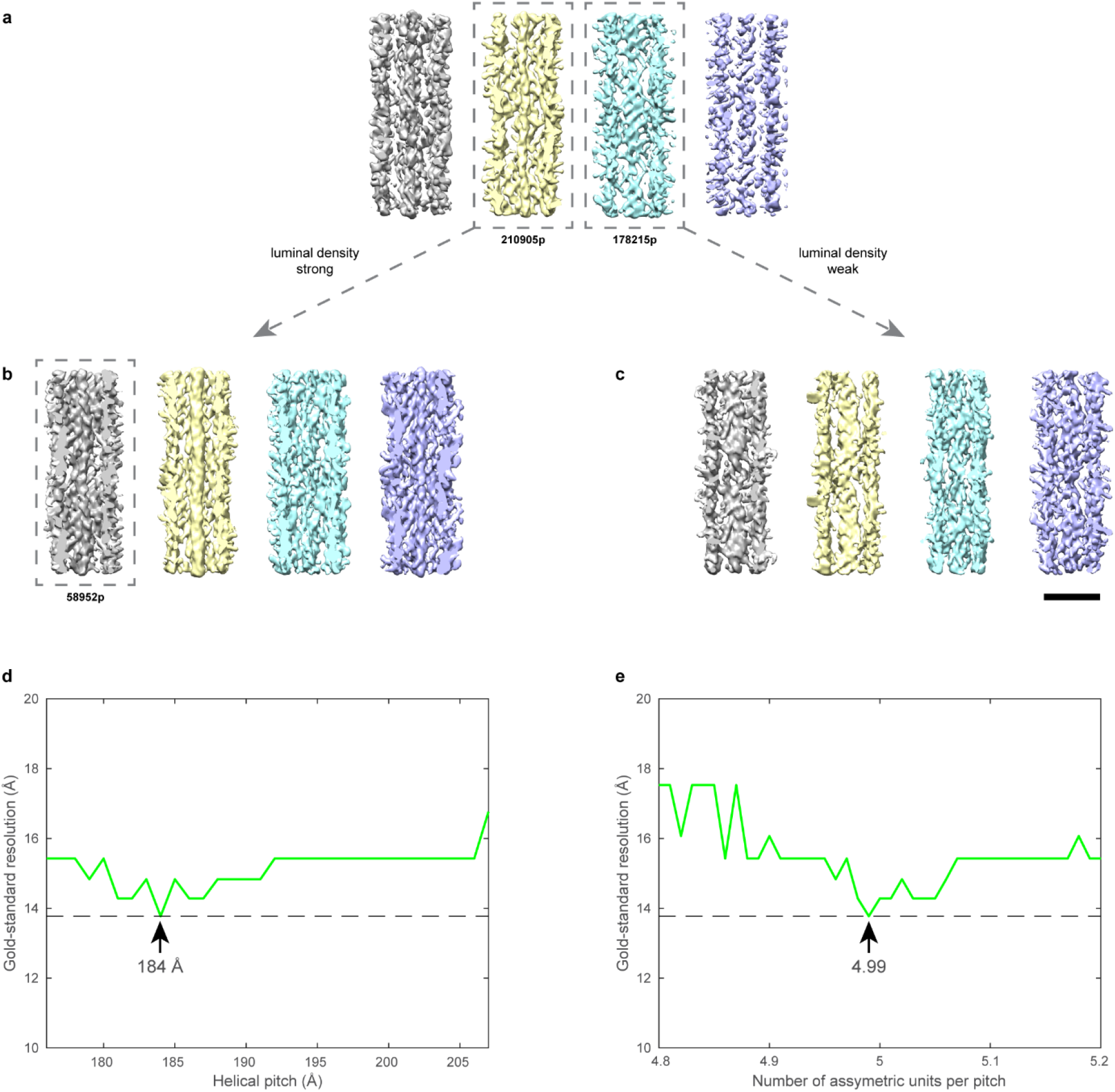
Helical 3D classification of VIF segments. (**a**) VIF segments (615,106 particles with a size of 38 × 38 nm^2^) were grouped by helical 3D classification in four classes (c1-c4). A helical symmetry with a helical pitch of 186 Å and a helical rise of 37 Å were assumed. The 3D class averages differ with respect to tube diameter and luminal density. Classes c1 and c2 show pronounced luminal densities, whereas c3 and c4 show weak or no luminal densities. This 3D classification defines which of the segments will be mapped as VIF state-1 (c1 and c2, in total 324,386 particles) or VIF state-2 (c3 and c4, in total 290,720 particles) on the assembled filaments (Fig. 2g). Classes c2 and c3 were further classified. The results are shown in (**b**) and (**c**), respectively. Scale bar 10 nm. (**d**) Those segments assigned to c1 in (**b**) (58,952 particles), were further analyzed by applying two more exhaustive helical parameter searches. The result of the first helical search is shown in (**d**). Here the number of asymmetric subunits in one pitch was kept constant at n=5.0, but the helical pitch was varied between 176 Å to 207 Å in 1 Å increments. This effectively translates into varying the helical rise h_r_ and keeping the helical twist h_t_ constant at 72°. The green curve shows the gold-standard resolution values ^49^, recorded with 3D Refine jobs in RELION ^48^. The global resolution optimum was found at a helical pitch of 184 Å (⇒ h_r_=36.8 Å, h_t_=72°). The result of the second helical search is shown in (**e**). Here we kept the helical pitch constant at 185.6 Å and varied n between 4.8 and 5.2. The optimal n was found at 4.99, therefore h_r_=37.2 Å and h_t_=72.2°. These helical parameters were used in the final 3D refinement as starting values for a local helical symmetry search as implemented in RELION.

**Supplementary Figure 7.**
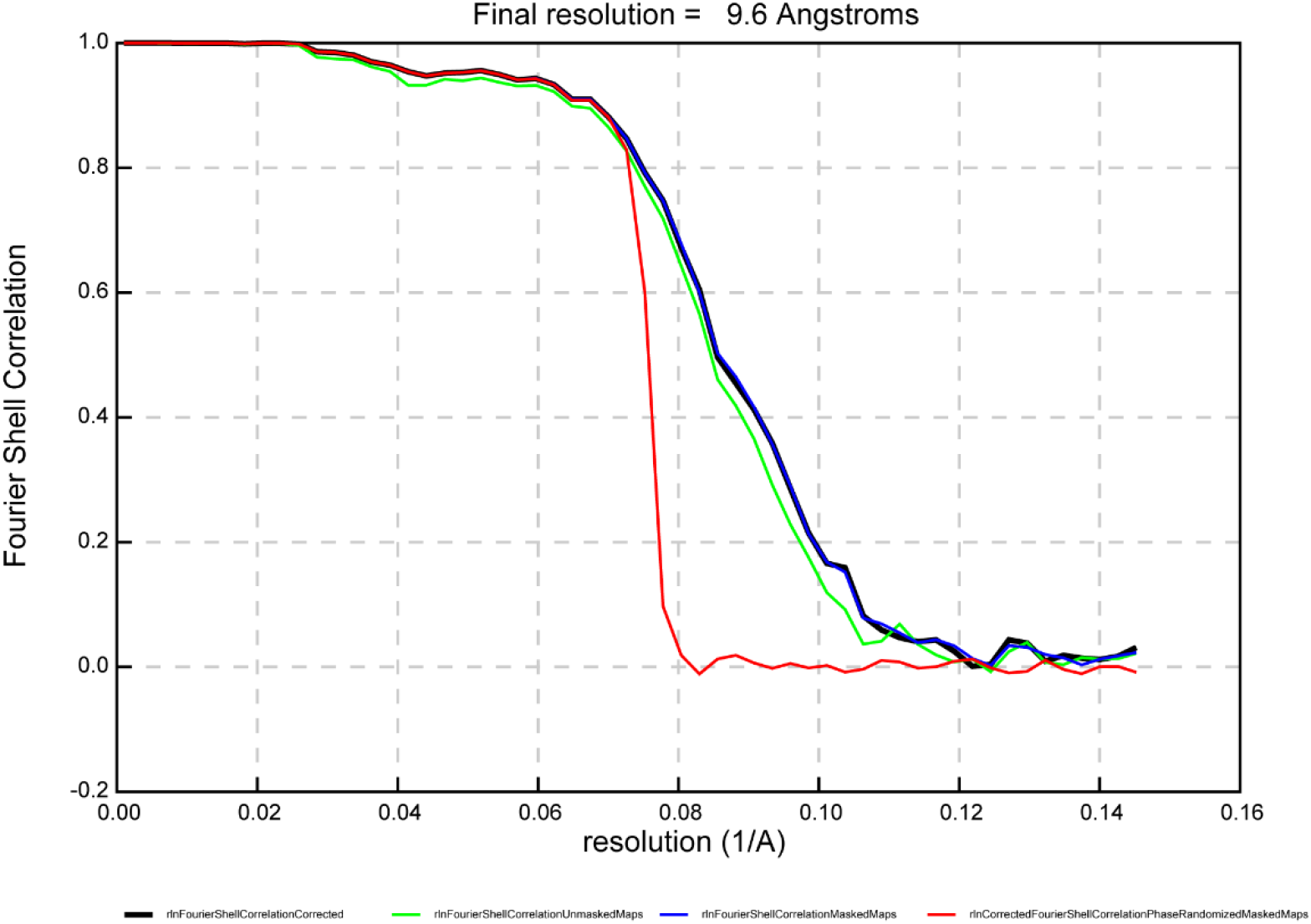
Resolution of the resulting VIF state-1 structure. The structure reached a resolution of 9.6 Å.

**Supplementary Figure 8.**
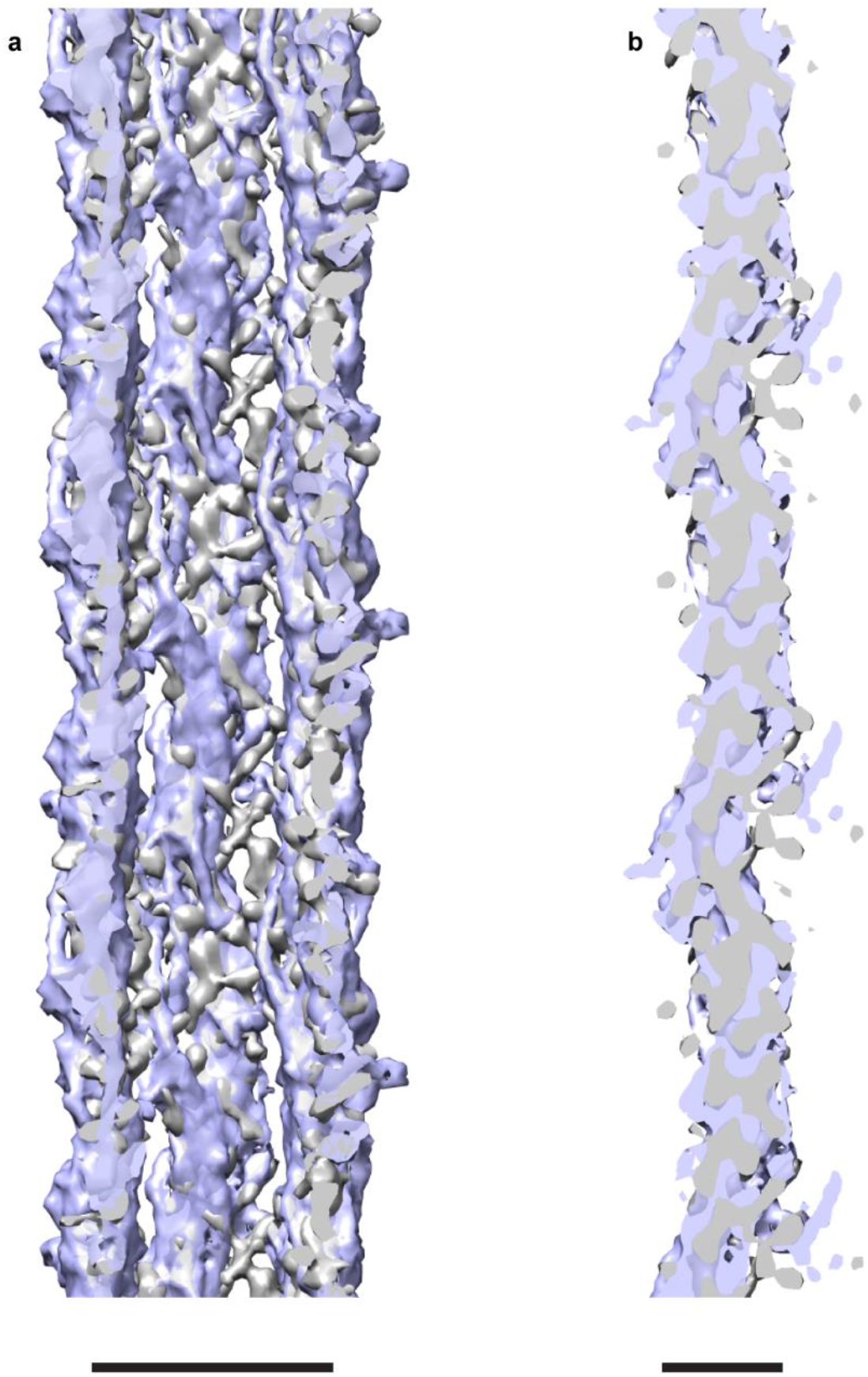
Docking of the VIF model. (**a**) The VIF model was converted into an electron density map (blue structure) and docked into the VIF structure (grey density). The correlation coefficient between the two structures is 0.81. Scale bar 10 nm. (**b**) The view sections through one of the protofibrils. The overall shape of the VIF model (blue density) matches the overall shape of the VIF structure (grey density). Scale bar 5 nm.

**Supplementary Figure 9.**
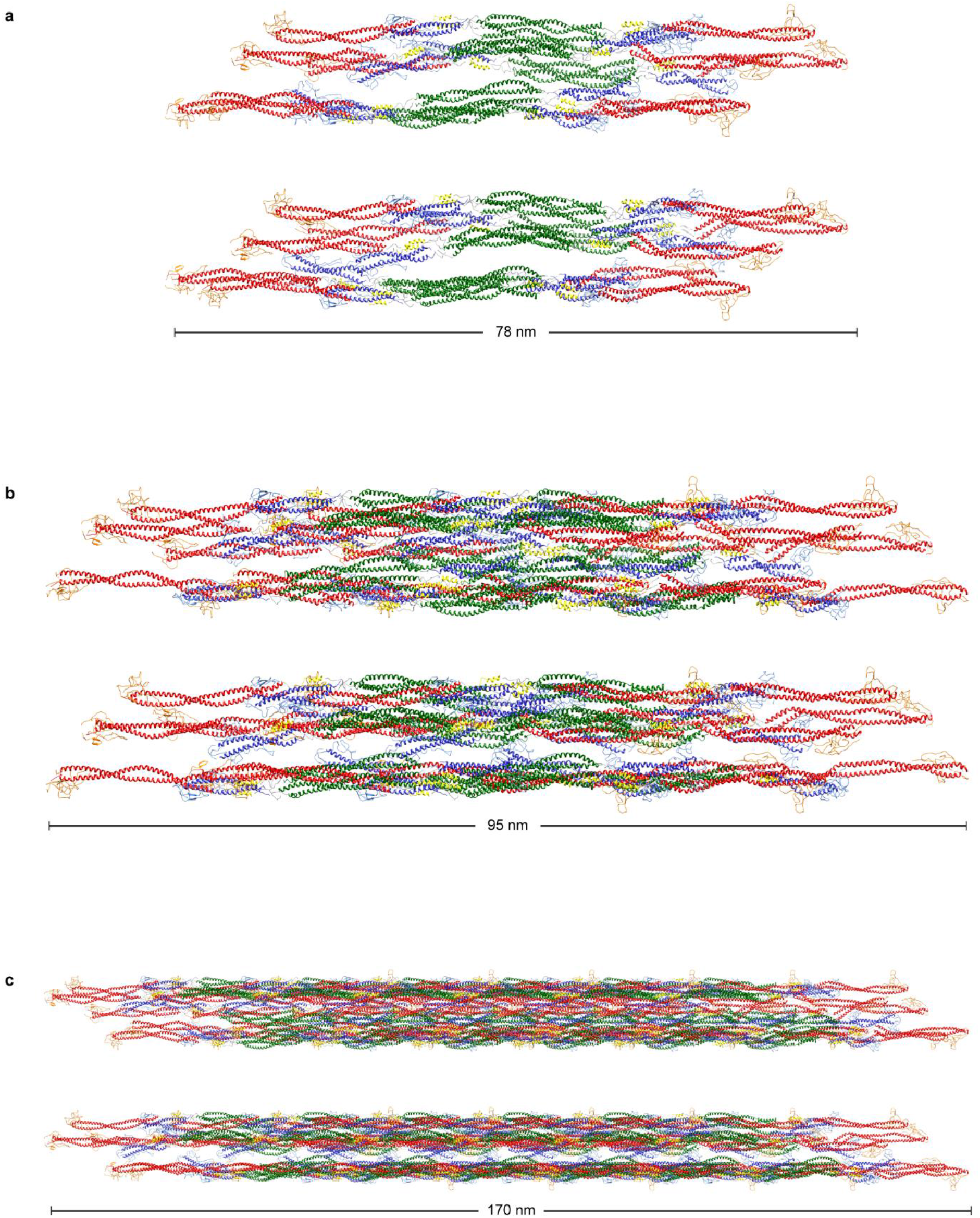
Gallery of VIF models. Three VIF models were constructed based on the measured helical symmetry and the known structure of the vimentin tetramer ^29^. The models differ in the number of tetramers: (**a**), 5 tetramers, (**b**), 10 tetramers, and (**c**), 30 tetramers, respectively. Each model is depicted in two views. The second view is rotated 18° around the filament axis compared to the first view.

**Supplementary Figure 10.**
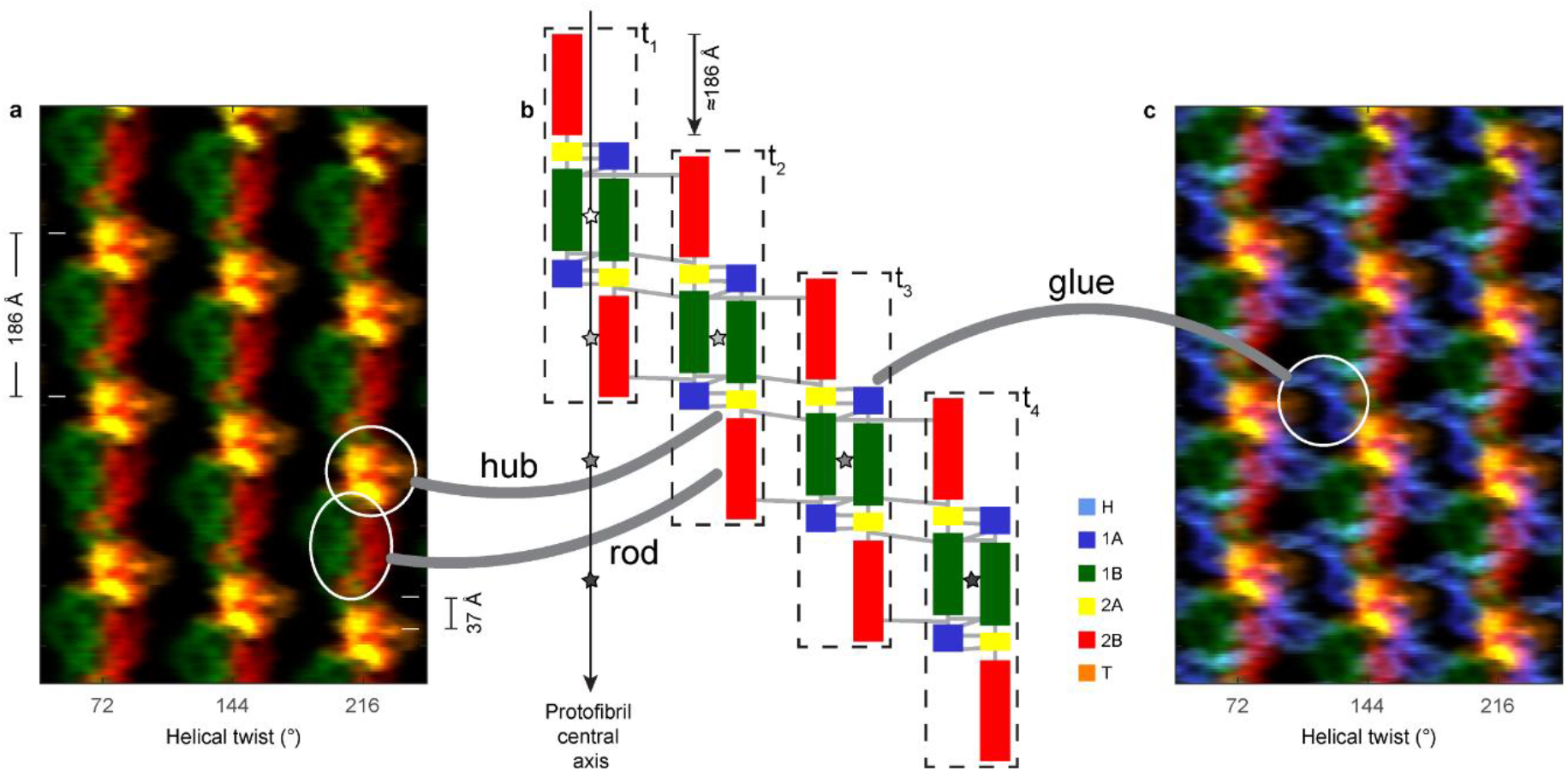
Molecular architecture of VIFs. (**a**) Illustration of the helical lattice of in-situ polymerized VIFs (three of five protofibrils ^32^ are depicted). The angular spacing between the protofibrils is ~72° (helical twist). Identical features along the same protofibril repeat every ~186 Å (helical pitch). Identical features between neighboring protofibrils are translated by ~37 Å (helical rise). The 1A and H domains are omitted in this image, so there are no connections visible between the protofibrils. (**b**) Illustration of the domain structure of a protofibril. Four A_11_ vimentin tetramers ^29,55^ are outlined with dashed rectangles (t_1_,…,t_4_) and their centers of mass are indicated with asterisks. A protofibril is constructed by successive translations of tetramers in steps of the helical pitch. Therefore, the mass centers of the tetramers are aligned along the protofibril central axis with a distance of the helical pitch length. In the illustration, we additionally added a progressive translation of the tetramers from left to right, in order to have an unobstructed view on the individual tetramers. Since the lateral stagger between two antiparallel dimers in the A_11_ vimentin tetramer is similar to the measured helical pitch, the tetramer with index i+1 along a protofibril creates an A_12_binding interface ^55^ with the tetramer with index i. Therefore, our VIF model agrees with the previously determined cross-links ^55^, which are represented by the domain connecting lines. (**c**) The 1A and H domains protrude into the space between the protofibrils to form the glue regions.

